# The rice bundle sheath produces reactive oxygen species during high light stress via NADPH oxidase

**DOI:** 10.1101/2020.07.06.189381

**Authors:** Haiyan Xiong, Lei Hua, Ivan Reyna-Llorens, Yi Shi, Kun-Ming Chen, Nicholas Smirnoff, Johannes Kromdijk, Julian M. Hibberd

## Abstract

When exposed to high light plants produce reactive oxygen species (ROS). In *Arabidopsis thaliana* local accumulation of ROS preferentially takes place in bundle sheath strands, but little is known about how this response takes place. Using rice and the ROS probes diaminobenzidine and 2’,7’-dichlorodihydrofluorescein diacetate, we found that after exposure to high light, ROS were produced more rapidly in bundle sheath strands than mesophyll cells. This response was not affected either by CO_2_ supply or photorespiration. Consistent with these findings, deep sequencing of mRNA isolated from mesophyll or bundle sheath strands indicated balanced accumulation of transcripts encoding all major components of the photosynthetic apparatus. However, transcripts encoding several isoforms of the superoxide/H_2_O_2_-producing enzyme NADPH oxidase were more abundant in bundle sheath strands than mesophyll cells. ROS production in bundle sheath strands was reduced by blocking NADPH oxidase activity pharmacologically, but increased when the bundle sheath preferential *RBOHA* isoform of NADPH oxidase was over-expressed. NADPH oxidase mediated accumulation of ROS in the rice bundle sheath was detected in etiolated leaves lacking chlorophyll indicating that high light and NADPH oxidase-dependent ROS production is not dependent on photosynthesis.

## Introduction

Under high-light conditions the capacity for light capture during photosynthesis can exceed use. This can lead to damage, generate signals promoting repair, and also initiate responses allowing acclimation (Asada, 1999; Li et al., 2009; Mullineaux et al., 2018). One source of damage is an increase in the production of reactive oxygen species (ROS). For example, oxygen photoreduction, largely at Photosystem I can result in superoxide and hydrogen peroxide. The other source is singlet oxygen which is formed by interaction of oxygen with triplet state chlorophyll in Photosystem II (Noctor and Foyer, 1998; Li et al., 2009; Murchie and Niyogi, 2011). As ROS are potentially harmful, with the ability to damage Fe-S proteins, oxidise amino acid residues and generate further radicals and reactive electrophiles resulting in lipid peroxidation and DNA damage, photosynthetic organisms have evolved a variety of mechanisms to minimize over-excitation of the photosystems. These range from transcriptional responses mediated by retrograde signaling between chloroplast and nucleus (Rossel et al., 2002; Kimura et al., 2003; Kleine et al., 2007; Li et al., 2009; Ruckle et al., 2012; Vogel et al., 2014; Dietz, 2015; Crisp et al., 2017; Mullineaux et al., 2018) to more immediate remodeling of light harvesting structures to dissipate excess excitation energy (Muller, 2001; Ruban, 2016).

Processes that dissipate energy in excess of that used by the photosynthetic electron transport chain are collectively known non-photochemical quenching (NPQ) mechanisms, and their induction is thought to reduce damage to the photosynthetic apparatus caused by synthesis of ROS (Demmig-Adams and Adams, 1992; Muller, 2001). As such, the scavenging/antioxidant network to remove ROS and repair damge is complex (Asada, 1999; Mullineaux et al., 2006; Miller et al., 2010). Notably, although in C_3_ species such as *Arabidopsis thaliana*, mesophyll cells contain the majority of chlorophyll in a leaf, after exposure to excess light ROS accumulate preferentially in bundle sheath cells that surround veins (Fryer et al., 2002; Fryer et al., 2003; Mullineaux et al., 2006; Galvez-Valdivieso et al., 2009). ROS have been implicated in rapid systemic signaling responses initiated after various abiotic and biotic stresses including heat, wounding and pathogen attack (Fichman and Mittler, 2020). Such ROS mediated signaling from a locally perturbed leaf can lead to stomatal aperture being altered in distant leaves, is associated with the hormones abscisic and jasmonic acid, and dependent on the plasma membrane localized NADPH oxidase (Respiratory Burst Oxidase Homolog D (RBOHD) (Devireddy et al., 2018; Devireddy et al., 2020).

In contrast to C_4_ species in which the role of the bundle sheath in fixing CO_2_ by RuBisCO has been understood for decades (Hatch, 1987), this cell-type is poorly characterized in C_3_ species. Whilst the bundle sheath of C_3_ plants contains chloroplasts that accumulate starch (Miyake and Maeda, 1976; Kinsman and Pyke, 1998) they are not as numerous as those in the mesophyll and reducing chlorophyll accumulation in these cells has limited impact on photosynthesis (Janacek et al., 2009). Rather, in *A. thaliana* the bundle sheath is thought to be specialized in sulphur metabolism, glucosinolate biosynthesis (Gigolashvili et al., 2007; Koroleva et al., 2010; Aubry et al., 2014) and transport of water and solutes in and out of the leaf (Aubry et al., 2014). In particular, stress responsive activation of aquaporins in bundle-sheath cells are important in hydraulic conductivity of the whole leaf (Shatil-Cohen et al., 2011; Sade et al., 2014; Sadey et al., 2015). Consistent with this, bundle sheath cells more generally have been proposed to play a role in maintaining hydraulic integrity of the xylem (Sage, 2001; Griffiths et al., 2013) and in regulating flux of metabolites in and out of the leaf (Leegood, 2008).

To our knowledge none of these previous studies explain how the bundle sheath of C_3_ plants preferentially accumulates ROS. One possibility is that supply of atmospheric CO_2_ to cells around the veins is limited and so inorganic carbon present in the transpiration stream provides CO_2_ to photosynthesis (Hibberd and Quick, 2002; Brown et al., 2010). If this were the case, when stomata close, provision of CO_2_ from veins could slow activity of the Calvin Bassham Benson cycle compared with chlorophyll de-excitation in the light harvesting complexes (Galvez-Valdivieso et al., 2009). Although, proximity of bundle sheath cells to veins could provide an efficient mechanism to initiate systemic acclimation to high-light stress (Mullineaux et al., 2006) the mechanism/s by which ROS accumulate in bundle sheath cells is unclear, and to our knowledge how common this response is beyond *A. thaliana* is not known.

Using rice, we show that ability of the C_3_ bundle sheath to preferentially accumulate ROS in response to high light is found in the monocotyledons as well as the dicotyledons. We found no evidence that ROS accumulation in the C_3_ bundle sheath was due to limited CO_2_ supply, nor to production of H_2_O_2_ from photorespiration – in fact we detected clear ROS accumulation in bundle sheath cells of C_4_ species in which photorespiration is essentially abolished. We also found no evidence for an imbalance between transcript abundance of genes encoding components of light harvesting apparatus and the Calvin Bassham Benson cycle. However, we did find that transcripts encoding NADPH oxidases accumulate preferentially in bundle sheath cells. Pharmacological treatment to block their activity and overexpression reduced and increased accumulation of ROS in the bundle sheath of rice respectively. Although accumulation of ROS in the bundle sheath was strongest in green leaves containing light harvesting apparatus, accumulation was still detected in etiolated leaves.

## Results

### Veins and bundle sheath cells of rice preferentially accumulate reactive oxygen species in response to high light

Rice leaves were exposed for 90 minutes to a light intensity tenfold higher than that used for growth. As expected, this led to a rapid and sustained reduction in the chlorophyll fluorescence parameters Fv’/Fm’ and Fq’/Fm’ (Figure 1A&B) that report the maximum efficiency of Photosystem II and its operating efficiency respectively. Over the same period, photochemical quenching first decreased and then recovered slowly, whilst non-photochemical quenching increased steadily (Figure 1C). Representative images of Fv’/Fm’ over this time-course indicated that responses of the photosynthetic apparatus to high light were relatively homogenous across the leaf (Figure 1D). Together, these data show that subjecting rice leaves to excess light led to the expected response of Photosystem II efficiency.

**Figure 1.**
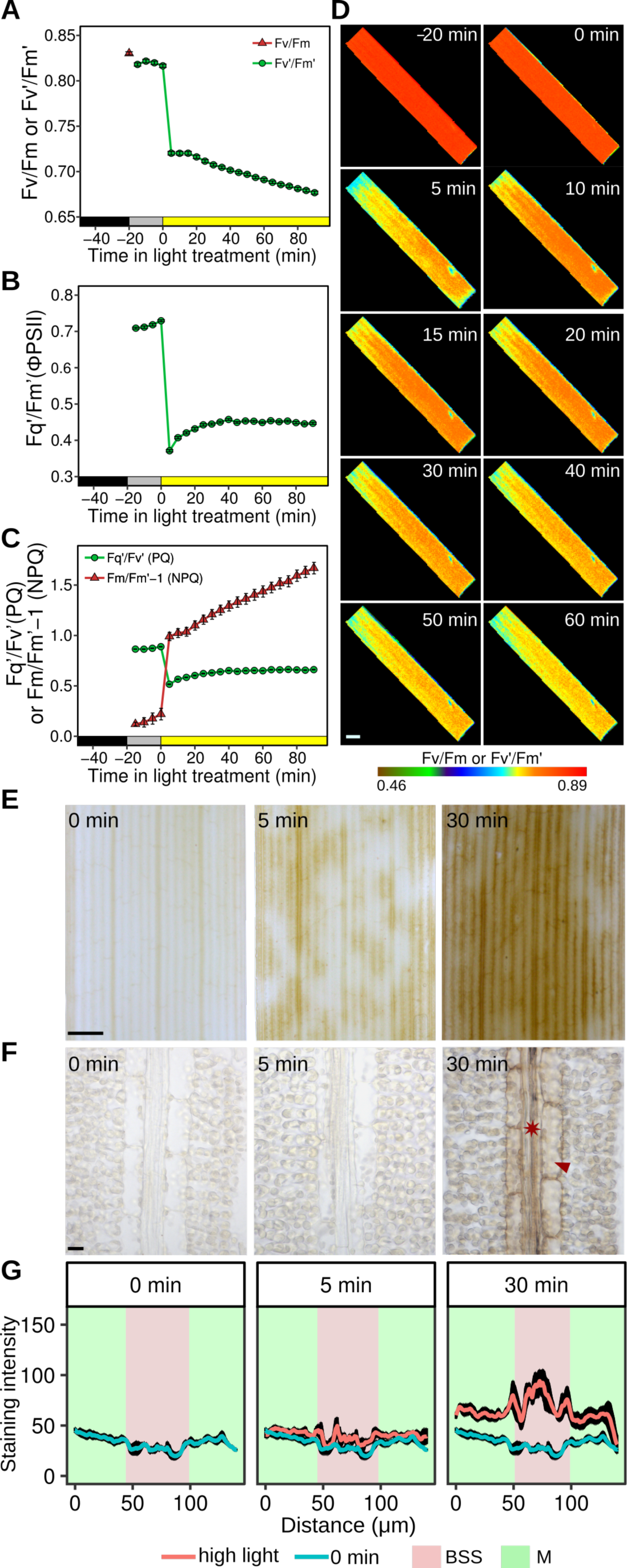
Rice bundle sheath strands preferentially accumulate the DAB polymerization product in response to high light. (A-C) Chlorophyll fluorescence parameters associated with dark-adapted leaves being moved into the light intensity of growth for 20 min, and then moved to a 10-fold higher intensity of light. (A) Dark-adapted F_v_/F_m_ and F_v_’/F_m_’ (B) Quantum efficiency of PSII (F_q_’/F_m_’ or FPSII) and (C) Photochemical (PQ) and Non Photochemical Quenching (NPQ). Data shown represent mean and standard error from 16 leaves. (D) Representative images from the chlorophyll fluorescence imager showing responses were reasonably homogenous across the leaf. Scale bar = 2 mm. (E) High light stress led to strong staining from the DAB polymerization product in bundle sheath strands arranged along the proximal to distal axis of the leaf blade. After five minutes of high light staining is evident, but at thirty minutes it is stronger and more homogenous in these bundle sheath strands. Scale bar = 1 mm. (F) Representative image from paradermal sections show that cells accumulating DAB stain are veins (asterisk) and bundle sheath cells (arrowhead). Scale bar = 10 μm. (G) Quantitation of DAB stain in mesophyll and bundle sheath strands. Data are presented as mean (red or blue line) and one standard error from the mean, n= 4).

We next tested whether preferential accumulation of ROS in bundle sheath cells as reported in *A. thaliana* (Fryer et al., 2002; Fryer et al., 2003; Galvez-Valdivieso et al., 2009) was detectable in rice. The cytochemical dye 3, 3-diaminobenzidine (DAB) is routinely used to detect H_2_O_2_. It reacts with H_2_O_2_ to form a brown polymerization product, the reaction being accelerated by peroxidase (Thordal-Christensen et al., 1997). In contrast to the relatively homogenous alterations to chlorophyll fluorescence parameters reporting on the activity of Photosystem II (Figure 1D), preferential accumulation of the DAB polymerization product (hereafter referred to as DAB) was detected in patches of longitudinal files of cells after five minutes, and then in almost all files of these cells after thirty minutes of exposure to high light (Figure 1E). Paradermal sections from leaves were generated in order to determine the specific cell types involved, and this showed that thirty minutes after exposure to high light the strongest DAB signal was associated with veins and bundle sheath cells (Figure 1F). We refer to these cells as bundle sheath strands (BSS) as they include both the bundle sheath and the vascular strands. Quantitation of this signal from multiple sections confirmed that bundle sheath strands consistently accumulated more DAB than the surrounding mesophyll cells (Figure 1G) and more dense sampling indicated that the increase in DAB was first detectable ten minutes after the transfer to high light (Figure S1A&B). From around fifteen minutes after the treatment, DAB also increased in mesophyll cells, but the signal was always stronger in bundle sheath strands (Figure S1A&B). We used the ROS sensitive fluorescent dye 2’,7’-dichlorodihydrofluorescein diacetate (H_2_DCFDA) to provide independent evidence that rice bundle sheath strands were particularly responsive to high light. In the presence of peroxidases and radicals generated by ROS in living plant cells, H_2_DCFDA produces highly fluorescent 2’,7’-dichlorofluorescein (DCF) (Winterbourn, 2014). Consistent with the results obtained with DAB, when H_2_DCFDA was supplied to rice leaves, high light led to brighter green fluorescence in bundle sheath strands than neighbouring cells (Figure S2A). To exclude the possibility that H_2_DCFDA had not uniformly penetrated mesophyll cells, we supplied a subset of leaves with exogenous H_2_O_2_ (Figure S2B). This led to DCF fluorescence from both mesophyll as well as bundle sheath strands indicating that the increased signal in the bundle sheath after exposure to high light was unlikely to be an artefact of incomplete transport into all cells of the leaf (Figure S2B). Localised staining also rules out direct dye oxidation by light (Winterbourn, 2014). We conclude that as with *A. thaliana*, veins and bundle sheath cells of rice preferentially accumulate ROS in response to high light treatment and thus that this may be a property found in both dicotyledons and monocotyledons.

### Over-capacity in light harvesting compared with Calvin-Benson-Bassham cycle capacity is unlikely the cause of DAB accumulation in rice bundle sheath strands

Excess excitation energy that cannot be fully utilized by carbon assimilation and other metabolic processes in chloroplasts gives rise to ROS during exposure to high light intensity (Fryer et al., 2003; Apel and Hirt, 2004; Bechtold et al., 2008). The greater production of ROS in the BSS compared with mesophyll cells implies a greater capacity for ROS production or a limitation imposed by CO_2_ assimilation rate. As bundle sheath strands are distant from stomata, and in contact with fewer intercellular air spaces than mesophyll cells, we reasoned that they may be CO_2_ limited. Thus, flux through the Calvin-Benson-Bassham cycle may be constrained compared with activity of the photosynthetic electron transport chain. If this were the case, reducing and increasing the CO_2_ concentration around leaves would be expected to respectively enhance and repress preferential DAB staining in bundle sheath strands. However, we found evidence for neither. High light exposure at 200 ppm [CO_2_], which would restrict activity of the Calvin Benson Bassham cycle, led to preferential but similar DAB staining in the bundle sheath strands and 2000 ppm [CO_2_], which should saturate RuBisCO in bundle sheath as well as mesophyll cells, failed to abolish the preferential DAB staining in these cells (Figure 2A&B).

**Figure 2.**
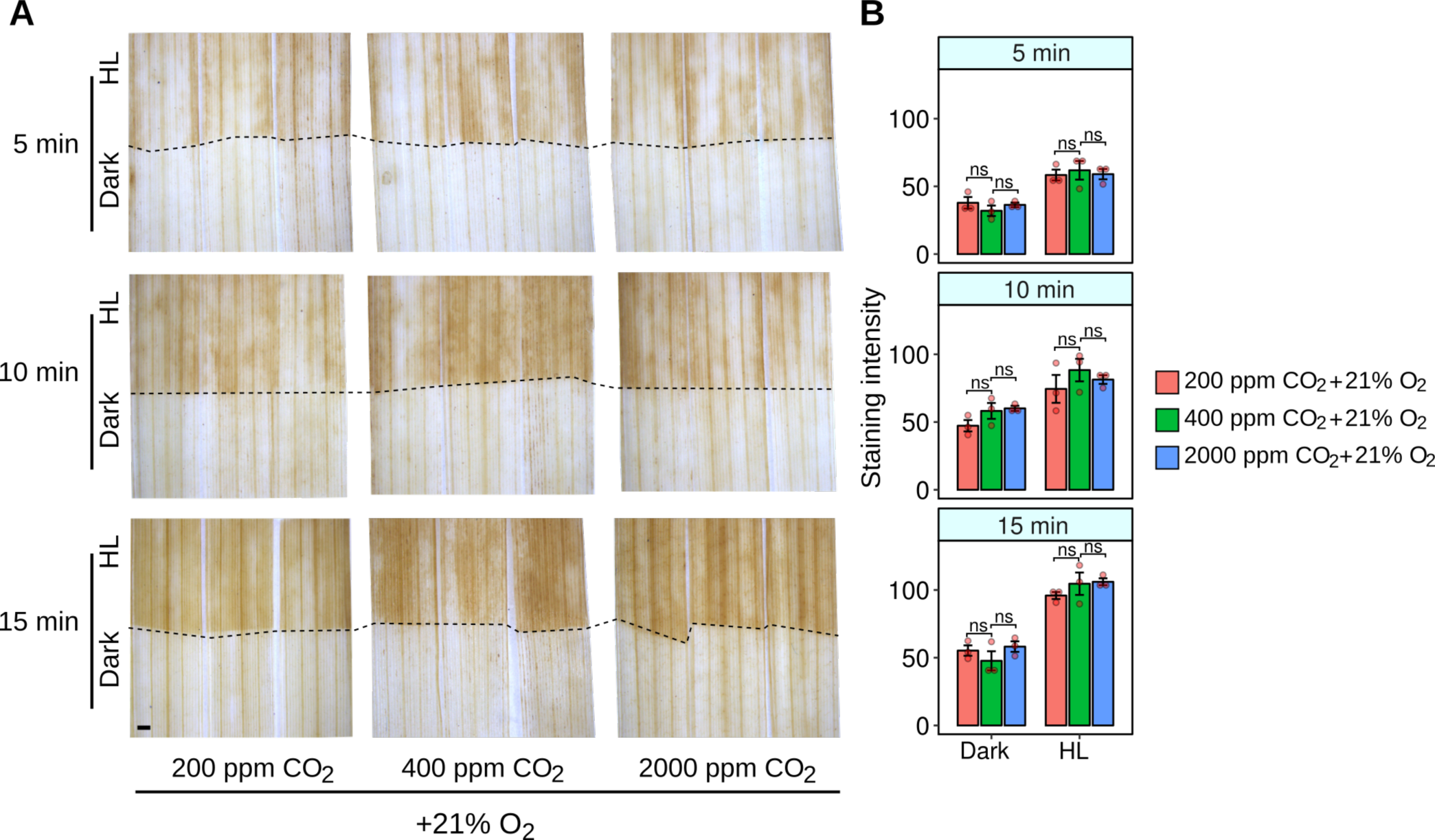
Altering CO_2_ supply has little effect on the high light response of the rice bundle sheath. (A) Neither reducing (200 ppm) nor increasing (2000 ppm) CO_2_ had a clear effect on DAB staining at 21% oxygen. (B) Quantitation of DAB staining of leaves subjected to low or high CO_2_. ANOVA indicated no significant statistical difference associated with CO_2_ treatment (p=0.22). Scale bar represents 1 mm.

The oxygenation reaction of RuBisCO requires the photorespiratory pathway to detoxify the initial product phosphoglycolate, and in so doing H_2_O_2_ is released by glycolate oxidase in peroxisomes. It is therefore possible that rapid DAB staining in bundle sheath strands is due to high rates of photorespiration in this cell type. To test this, we reduced the oxygen tension to 2%, but found that this had no clear effect on DAB staining compared with controls (Figure 3A&B). Moreover, when leaves of the C_4_ species *Gynandropsis gynandra, Setaria italica, Zea mays* and *Sorghum bicolor*, which generate up to tenfold higher concentrations of CO_2_ in bundle sheath cells and therefore minimal activities of photorespiration were exposed to high light, DAB still accumulated in these cells over a similar time-course (Figure 3C). It therefore appears unlikely that preferential DAB staining in either C_3_ or C_4_ bundle sheath cells is caused by H_2_O_2_ produced during photorespiration.

**Figure 3.**
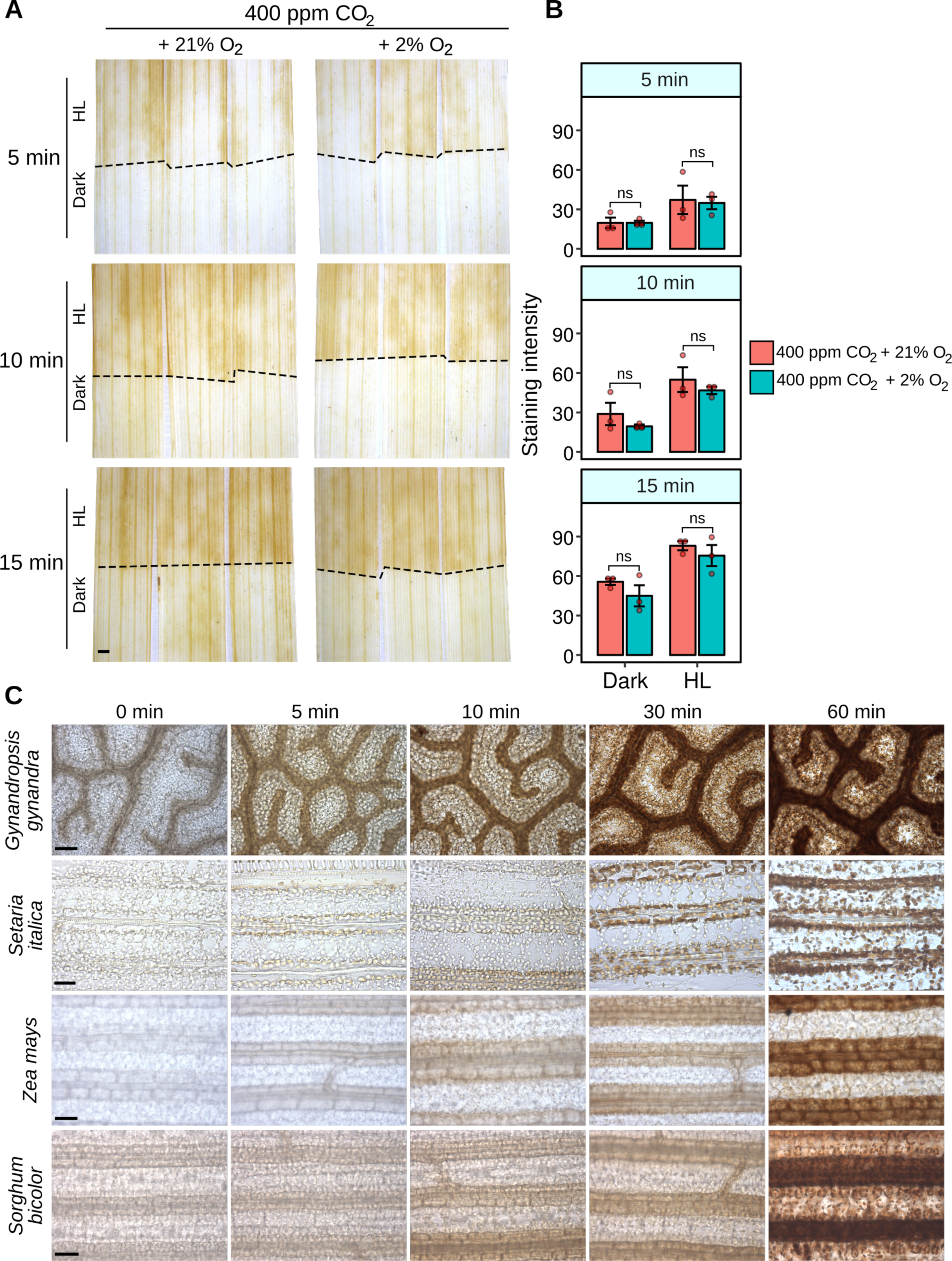
Photorespiration is not the source of the high light dependent accumulation of DAB stain in bundle sheath strands. (A) Representative images of DAB staining leaves exposed to 750 μmol m^-2^ sec^-1^ photon flux density for 5, 10 or 15 mins in 400 ppm CO_2_ and either 21% or 2% O_2_. Photorespiration is limited by 2% O_**2**_. (B) Quantification of DAB staining. ANOVA indicated no significant statistical difference associated with O_2_ treatment (p=0.0933). (C) DAB staining leaves of C_4_ plants *Gynandropsis gynandra, Setaria italica, Zea mays, Sorghum bicolor* exposed to high light. Although C_4_ plants have limited photorespiration in the bundle sheath, DAB staining was still detected in this tissue. Scale bars represent 1 mm (A) and 50 μm (B).

Taken together our results imply that accumulation of ROS in the bundle sheath of C_3_ and C_4_ plants is unlikely to be caused by limited capacity of the Calvin-Benson-Bassham cycle compared with ability to harvest light energy. Furthermore, the data obtained by suppressing photorespiration either transiently in C_3_ leaves, or more permanently in C_4_ leaves, are inconsistent with the notion that photorespiratory derived H_2_O_2_ is responsible for the rapid accumulation of ROS in bundle sheath cells.

### Transcriptome profiling indicates balanced expression of photosynthesis genes but elevated expression of genes encoding enzymes responsible for synthesis of reactive oxygen species in bundle sheath strands

To better understand the molecular basis for preferential accumulation of ROS in rice bundle sheath strands we carried out RNA-SEQ on this tissue. Laser capture microdissection was used to obtain mRNA from three biological replicates of bundle sheath strands or mesophyll cells derived from leaves that had not received a high light treatment (Figure 4A). Electropherograms showed that the RNA obtained was of good quality (Figure 4B). In total, over 165 million reads were generated using the Illumina sequencing platform. For each replicate, on average about 78.5% of reads were mapped uniquely to the Nipponbare reference genome (Supplemental Table 1). To enable comparison between samples, we normalized all read counts with DESeq2, and to reduce noise, poorly expressed genes with averaged normalized counts of < 10 in all samples were removed. This led to a total of 15,727 genes being identified as expressed. Among these, 15456 genes were expressed in both BSS and M cells, while 239 genes were only expressed in BSS and 32 genes only expressed in M cells (Figure 4C). Hierarchical clustering (Figure 4D) and principal component analysis (PCA; Figure 4E) showed strong clustering between biological replicates from each tissue. Indeed, the PCA showed that 92% of variance between replicates was associated with one major component that mapped onto the tissue from which RNA was isolated (Figure 4E). Using criteria of log_2_ fold change > 0.5 and an adjusted *P* value < 0.05, we defined 3170 genes as being more highly expressed in bundle sheath strands, and 2766 as more strongly expressed in mesophyll cells (Figure 4F).

**Figure 4.**
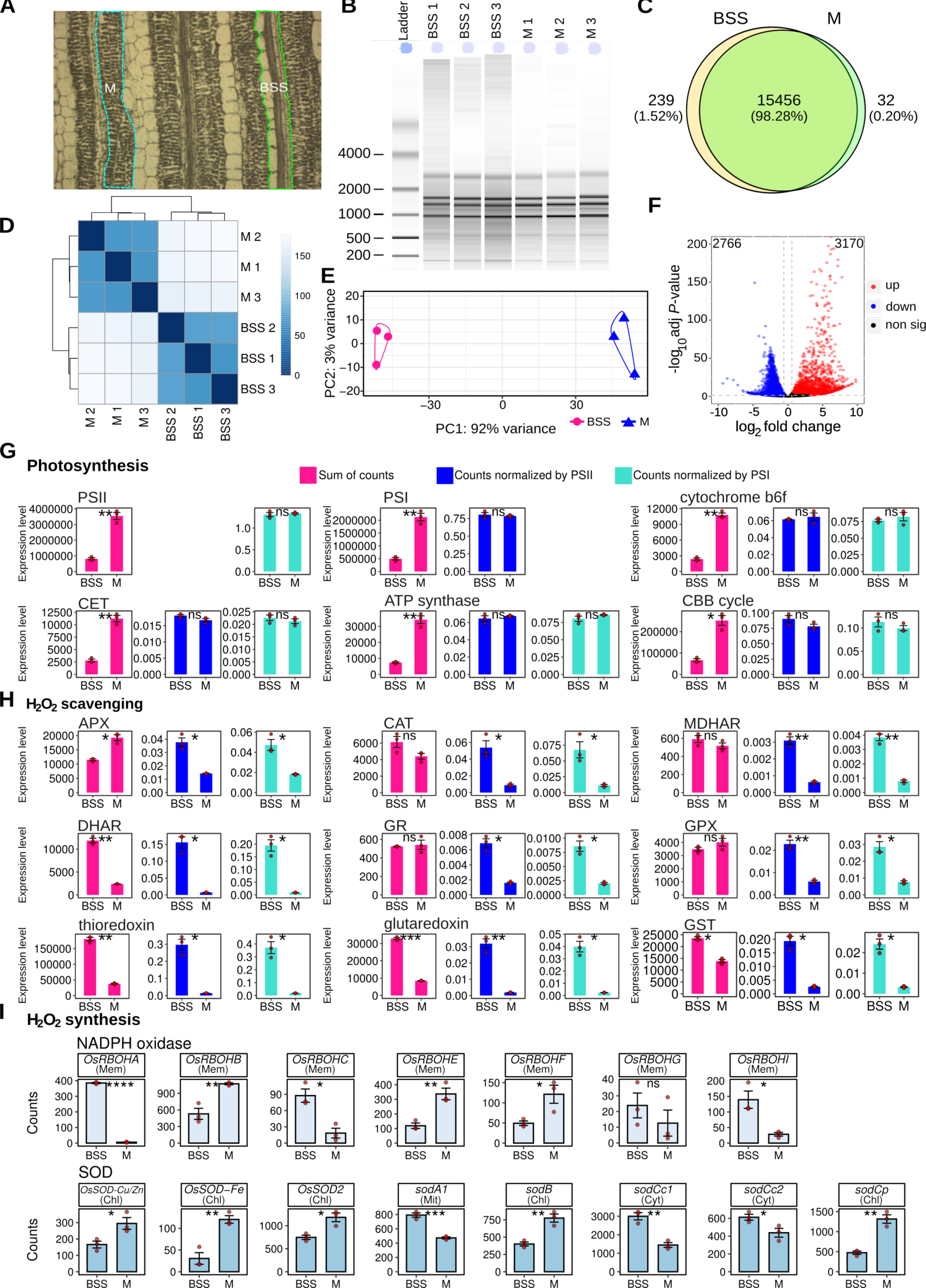
Global analysis mRNA from bundle sheath strands (BSS) and mesophyll (M) cells indicates balanced accumulation of transcripts encoding components of the photosynthetic electron transport chain and the Calvin Benson cycle, but that bundle sheath strands may have greater capacity to synthesise reactive oxygen species. (A) Bundle sheath strands (BSS) and mesophyll (M) were sampled for RNA using laser capture microdissection. (B) Electropherogram showing RNA quality. (C) Pie chart summarizing number of genes expressed in both BSS and M cells. (D) Hierarchical dendrogram and (E) Principal Component Analysis indicate that most variance was associated with tissue type. (F) Volcano plot showing number of differentially expressed genes between BSS and M. (G-I) Abundance of photosynthesis (G), H_2_O_2_ scavenging (H) and H_2_O_2_ synthesis (I) transcripts. For photosynthesis complexes the sum of all components of each complex is presented, and to take into account lower chloroplast content in bundle sheath strands these were normalized to either Photosystem I or II. Predicted subcellular localization of each NADPH oxidase (RBOH) and Superoxide dismutase (SOD) isoform is annotated (chl = chloroplast, mem = plasma membrane, mit = mitochondrial, cyt = cytoplasm.). T-tests indicate statistically significant differences (**** p< 0.0001, *** p< 0.001, ** p< 0.01, * p< 0.05).

We next analysed the abundance of transcripts derived from genes known to be associated with the production of ROS. Specifically, we focused on genes encoding the photosynthetic apparatus, and enzymes that either scavenge or synthesize ROS. In all cases, to provide an overview of these complex processes eigengene values were computed to take into account the fact that multiple genes encode these oligomeric protein complexes. Transcripts encoding components of Photosystem II, Photosystem I, the cytochrome *b*_*6*_*f* complex, cyclic electron transport and the ATP synthase were less abundant in bundle sheath strands compared with mesophyll cells (Figure 4G and Figure S3). However, the mesophyll contains a larger chloroplast compartment (Kinsman and Pyke, 1998) and so this is to be expected. We thus normalized transcript abundance for each complex to Photosystem II and Photosystem I (Figure S3). This showed that components of Photosystem II, the cytochrome b6f complex and Photosystem I accumulated stoichiometrically in both cell types (Figure 4G). In other words, there was no clear imbalance in transcripts encoding one part of the photosynthetic electron transport chain that might lead to impaired function. This was also true for transcripts encoding enzymes of the Calvin-Benson-Bassham cycle (Figure 4G) indicating that the relative capacities of the light-dependent reactions of photosynthesis and the Calvin-Benson-Bassham cycle are balanced similarly in bundle sheath strands and mesophyll cells. These data are consistent with our earlier finding that increasing the CO_2_ concentration around leaves failed to abolish accumulation of DAB staining in the rice bundle sheath (Figure 2A&B).

We also found no evidence from this analysis of transcript abundance that bundle sheath strands have lower ability to detoxify ROS than the mesophyll (Figure 4H). After normalization to transcripts for the Photosystems which would be an expected source of ROS during high light stress, transcripts encoding enzymes known to scavenge ROS were typically more abundant in bundle sheath strands compared with mesophyll cells (Figure 4H). However, transcripts encoding several NADPH-oxidase and superoxide dismutase proteins, which generate superoxide and H_2_O_2_ respectively were more abundant in bundle sheath strands compared with mesophyll cells (Figure 4I). In particular, transcripts encoding RBOHA, RBOHC and RBOHI were considerably more abundant in bundles sheath strands compared with mesophyll cells. Quantitative Polymerase Chain Reactions (Q-PCR) confirmed these findings (Figure S4). These data imply that the rapid increase in ROS in bundle sheath strands after exposure to high-light is unlikely associated with a limited ability to dissipate energy associated with the photosynthetic electron chain, but rather appears due to higher basal activities of proteins that synthesize ROS.

### NADPH-oxidase activity mediates the high light response in both green and etiolated leaves

To test whether NADPH-oxidase activity is important for ROS accumulation in bundle sheath strands of rice we used inhibitors to block its activity, as well as a previously reported overexpression line for the *OsRBOHA* gene that encodes the major RBOH isoform NADPH-oxidase A in rice (Wang et al., 2016). Two commonly employed inhibitors of flavin-linked enzymes, diphenyleneiodonium chloride (DPI) (O’Donnell et al., 1993) and imidazole (Iizuka et al., 1985), were used to assess the role of NADPH oxidase. Both inhibitors reduced ROS production in bundle sheath strands of rice after imposition of light stress (Figure 5A&B and Figure S5A&B). The incomplete repression of ROS production associated with DPI or imidazole treatment is consistent with some production being associated with photosynthesis-related processes. To test whether this was the case, we next exposed etiolated leaves that lack chlorophyll to high light in the absence or presence of each inhibitor. Despite these leaves lacking chlorophyll, DAB staining was detected after exposure to high-light (Figure 5C&D, Figure S5C&D). This is consistent with a significant amount of ROS being produced from non-photosynthetic pathways. Moreover, in leaves lacking chlorophyll, inhibition of NADPH-oxidase activity completely abolished ROS production in bundle sheath strands (Figure 5C&D, Figure S5C&D). Q-PCR on RNA isolated from etiolated leaves confirmed that transcripts derived from *NADPH-oxidase A, NADPH-oxidase* C and *NADPH-oxidase I* genes as well as superoxide dismutase genes were more abundant in bundle sheath strands than in mesophyll cells (Figure S6). Whilst a mutant allele for *NADPH-oxidase B*, which was preferentially expressed in mesophyll cells (Figure 4I) had no impact on DAB staining during high light treatment (Figure S7), leaves from an overexpressor of *NADPH-oxidase A* (Wang et al., 2016) showed increased DAB staining in rice bundle sheath strands compared with controls (Figure 5E&F and Figure S8). We thus conclude that *NADPH-oxidase A* is important in mediating the response to high light in bundle sheath strands of rice.

**Figure 5.**
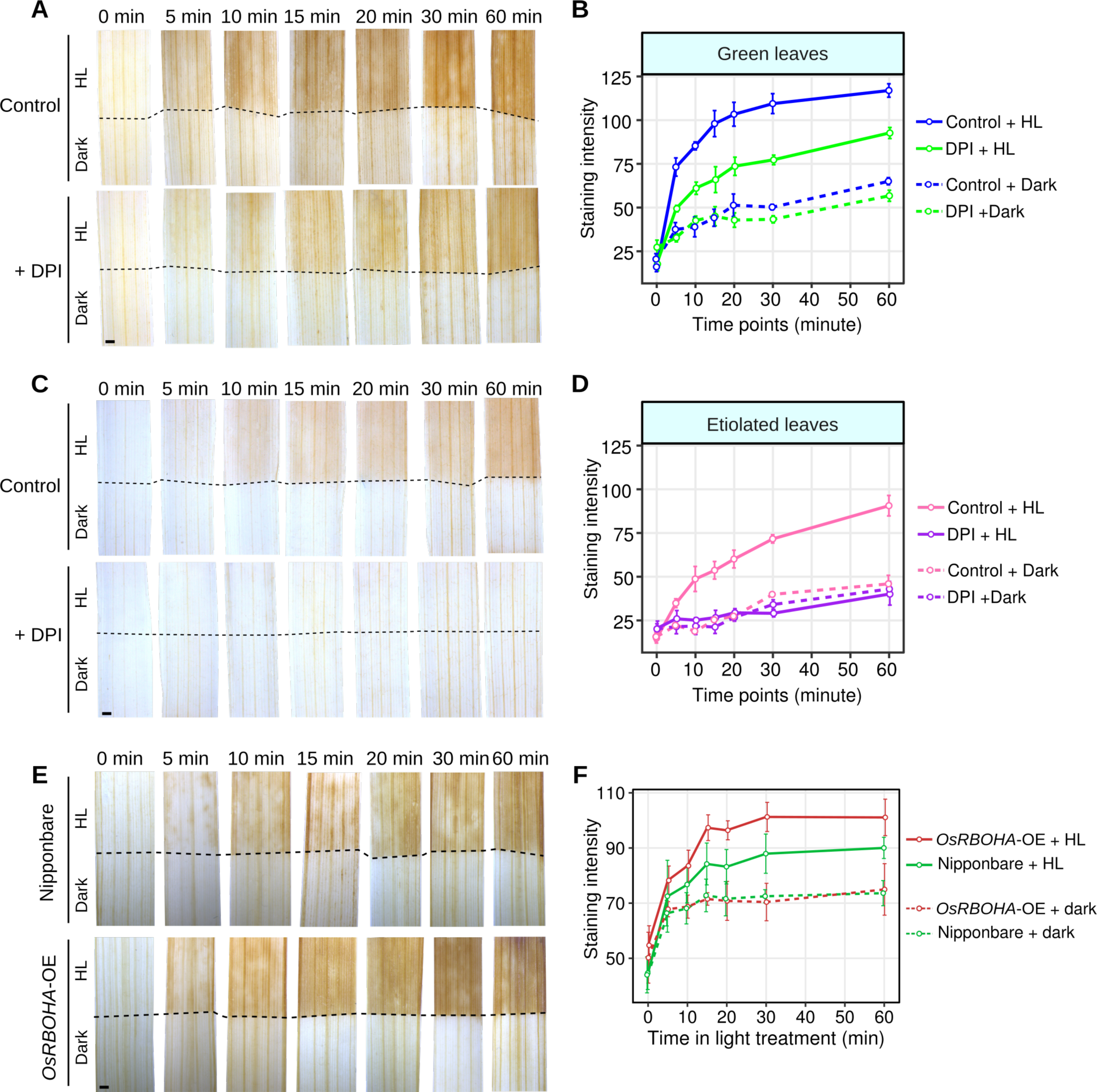
Inhibiting and increasing NADPH oxidase reduces and increases DAB staining respectively in bundle sheath strands of rice leaves. (A) Diphenyleneiodonium (DPI) a plasma membrane NAD(P)H oxidase inhibitor, partially inhibits DAB staining in bundle sheath strands of green leaves. (B) Quantitation of DAB staining over time in control compared with DPI and high light treated green leaves. (C) DPI completely inhibits DAB staining in etiolated leaves. (D) Quantitation of DAB staining over time in control and DPI high light treated etiolated leaves. (E) Overexpression of *OsRBOHA* increased DAB staining in bundle sheath strands after high light treatment. (F) Quantitation of DAB staining over time in green leaves from either the control or *OsRBOHA* overexpression line. In all cases the lower portion of the leaf (below the dotted line) was covered to provide an in-leaf control over the time-course. ANOVA showed DAB staining was reduced after DPI treatment in green leaves (p<0.001), etiolated leaves (p<0.001), and increased in *RBOHA* over-expressors (p<0.05). Scale bars represent 1 mm.

## Discussion

In the C_3_ species *A. thaliana* it has been known for some time that bundle sheath cells accumulate ROS in response to excess light (Fryer et al., 2002; Fryer et al., 2003; Mullineaux et al., 2006; Galvez-Valdivieso et al., 2009). The work presented here shows that a similar response is detected in bundle sheath strands of rice, a C_3_ species from the monocotyledons. Moreover, our analysis indicates that this phenomenon is not associated with C_3_ photosynthesis, but that it also takes place in bundle sheath strands from leaves of C_4_ plants. This was the case in C_4_ *Gynandropsis gynandra*, which is sister to the Brassicaceae, but also in the C_4_ grasses maize, sorghum and *Setaria*. These findings imply that the bundle sheath from dicotyledons and monocotyledons of both C_3_ and C_4_ species fulfills a role in sensing and responding to excess light.

To better understand the causes for production of ROS in bundle sheath strands we used rice. High light-induced ROS was not abolished by decreasing photorespiration in rice and could be seen in the bundle sheath strands of C_4_ species that lack photorespiration. Therefore, photorespiratory H_2_O_2_ production in peroxisomes *via* glycolate oxidase is not involved. The possibility that an imbalance between light harvesting and photosynthetic electron transport and use of excitation energy by the Calvin-Benson-Bassham cycle in bundle sheath cells was not supported by relative expression of genes encoding photosynthetic components. After normalization, these have similar relative expression in both cell types. This finding is consistent with analysis of the bundle sheath in C_3_ *A. thaliana* in which the photosynthetic apparatus is assembled and functional (Kinsman and Pyke, 1998; Janacek et al., 2009) despite lower levels of photosynthesis gene expression (Aubry et al., 2014). We also failed to collect any convincing physiological evidence that ROS production in the rice bundle sheath was associated with restricted activity of the Calvin-Benson-Bassham cycle as might be expected if provision of CO_2_ to this tissue distant from stomata was limiting. The accumulation of ROS was neither accentuated nor ameliorated when intercellular CO_2_ concentrations were reduced or increased respectively.

In contrast to a purely photosynthetic origin of ROS in bundle sheath strands, we have produced evidence that plasma membrane localized NADPH oxidase has a significant role. NADPH oxidase catalyses extracellular reduction of oxygen, forming superoxide in the apoplast. Membrane impermeable superoxide dismutates rapidly, producing H_2_O_2_ which is proposed to enter the cytosol through aquaporins. NADPH oxidase has numerous isoforms and is activated rapidly following numerous stimuli (Smirnoff and Arnaud, 2019). There is evidence for apoplastic ROS production in *A. thaliana* bundle sheath cells in response to high light (Galvez-Valdivieso et al., 2009) although NADPH depdendence was not established. We found that three NADPH oxidase isoforms (RBOHA, RBOHC and RBOHI) were more highly expressed in rice bundle sheath strands compared with mesophyll cells. Consistent with NADPH oxidase being involved, application of the flavoprotein inhibitors (DPI and imidazole) removed a large proportion of HL-induced ROS production. Moreover, a mutant allele and overexpressing line for *OsRBOHB*, which was more abundant in mesophyll cells showed no effect on ROS accumulation, whilst overexpression plants for *OsRBOHA* led to increased production of ROS in the bundle sheath. Thus, differential transcript abundance between the cell-types, pharmacological approaches to inhibit NADPH oxidase activity, and genetic perturbation of the *OsRBOHA* locus supported the notion that the bundle sheath strands have a greater ability than mesophyll cells to synthesize ROS in response to high light. Furthermore, the observation that etiolated leaves have high light induced ROS production provides key evidence that it is independent of photosynthesis and supports a role for NADPH oxidase. How NADPH oxidase is activated by light in etiolated leaves will need to be determined in the future. One possibility could be photoreceptor-mediated NADPH oxidase activation. Cryptochrome generates superoxide upon exposure to blue light and this has been proposed to contribute to part of its signaling role (El-Esawi et al., 2017). However, it is also possible that photoreceptors could activate NADPH oxidase. The results also suggest that part of the light-induced ROS derive from photosynthesis. Given that previous work has shown that chloroplasts release H_2_O_2_ in the light and that this could provide a high light signaling mechanism (Exposito-Rodriguez et al., 2017; Smirnoff and Arnaud, 2019), the possible differences in the function of chloroplast and NADPH oxidase-derived ROS requires investigation.

NADPH oxidase has previously been implicated in ROS responses induced by biotic and abiotic stresses. In response to a localized stress event, an NADPH oxidase-dependent ROS wave propagates between cells to initiate systemic responses in tissues distant from the original stress (Miller et al., 2010; Mittler et al., 2011; Zandalinas and Mittler, 2018; Fichman et al., 2019). Whilst we did not aim to investigate components underpinning such a wave, or downstream responses, our findings have some relevance to the role of NADPH oxidase in ROS production. The data imply that rice *RBOHA* plays a role in a local production of ROS in bundle sheath strands. How these findings relate to previous work on *A. thaliana* will require further work. The gene families encoding these proteins are large, and the phylogenetic distance between *A. thaliana* and rice means that direct orthologs are not always found, and function of these proteins may have diverged. However, *RBOHA* from rice, which appears to play an important role ROS accumulation in the bundle sheath in response to high light, is in the same orthogroup as *RBOHF* from *A. thaliana* (Figure S9A). Interestingly, compared with the whole leaf, transcripts of *AtRBOHF* were more abundant in bundle sheath cells (Figure S9B; (Aubry et al., 2014)). Thus, it is possible that these orthologs fulfill similar functions in the bundle sheath strands of both species. However, several lines of evidence point towards multiple pathways that both synthesize and propagate ROS production in response to stresses. For example, analysis of *A. thaliana* reported production of ROS in bundle sheath strands in leaves subjected to high light stress (Fryer et al., 2002; Fryer et al., 2003; Mullineaux et al., 2006; Galvez-Valdivieso et al., 2009), whilst in distant leaves ROS production is preferentially detected in mesophyll cells (Fichman et al., 2019; Fichman and Mittler, 2020). Moreover, differential changes in stomatal aperture of leaves subjected to stress compared with distant non-stressed leaves support a model in which two different signals are involved in systemic stomatal responses. One that is ABA based and associated with the vascular system (Schachtman and Goodger, 2008; KangasjÄrvi et al., 2009; Gorecka et al., 2014; Yoshida and Fernie, 2018) and another that uses ROS and travels through the plant (Devireddy et al., 2020). In fact, after a dark to light transition in *A. thaliana*, local changes to stomatal aperture were not dependent on *RBOHD*, but those changes associated with the systemic response were. *RBOHD* from *A. thaliana* is orthologous to *RBOHI* from rice and in both species transcripts appear to be more abundant in bundle sheath strands (Figure S9C&D and (Aubry et al., 2014)). This finding implies that the role of these two proteins may have been conserved since they diverged from the last common ancestor of rice and *A. thaliana*. Thus, downstream effects of production of ROS in the bundle sheath will require additional experimentation. One important advance in this area has been the ability to non-invasively monitor ROS either with reporters such as Hyper2 and roGFP (Exposito-Rodriguez et al., 2017; Nietzel et al., 2019), or with addition of fluorescent dyes to intact leaves to allow time-course analysis in locally stressed as well as distantly responding tissues (Fichman et al., 2019). Being able to apply these non-invasive approaches to tissues such as the bundle sheath that are deep in the leaf and therefore challenging to image will be informative in the future.

## Materials and methods

### Plant materials and high light stress

Rice (*Oryza sativa*. L) plants were grown for two weeks in compost in a growth chamber with a 16 h photoperiod with a photosynthetic photon flux density (PPFD) of 75 μmol m^-2^ s^-1^, day night temperatures of 28 and 26°C respectively, and relative humility of 60%. In all experiments, unless otherwise stated, IR64 was used. *OsRBOHA* and *OsRBOHB* overexpression lines and *Osrbohb* mutants as well as their background (Nipponbare), were grown under the same conditions. Etiolated plants were obtained by growing the plants in compost in the dark at 28/26°C for 2 weeks.

In all cases, the middle part (∼2.5 cm) of the second leaf was taken and infiltrated with dye solution (see below). Leaves were then subjected to 750 μmol m^-2^ s^-1^, which represented a tenfold increase above that of growth. High light was provided using a Clark-type oxygen electrode (LD2/3 oxygen electrode chamber) connected to an Oxylab control unit (Hansatech Instruments Ltd., Norfolk, UK) and temperature maintained at 28°C using a water tank. The top half of the leaves were illuminated by a liquid electronic display (LED) light source (Hansatech LH36-2), whilst the bottom half of the leaf was covered with tin foil to keep it in the dark.

### Chlorophyll fluorescence imaging

Chlorophyll fluorescence measurements were performed using a chlorophyll fluorescent imaging system (CF Imager, Technologica Ltd, Colchester, UK). ∼ 2 cm leaf strips of the middle part of second leaves were detached and floated on water in a 25-well square dish and transferred immediately into the CF Imager. The application of pre-programmed regimes of actinic growth light exposure times, dark periods, saturating light pulses, and the calculation and imaging of the parameters F_v_/F_m_, F_q_’/F_m_’, F_v_’/F_m_’, and NPQ (Baker, 2008) were performed using the manufacturers software-FluorImager.

### Detection of reactive oxygen species

H_2_O_2_ production in the cells was detected by staining with 3,3’-diaminobenzidine tetrachloride (DAB) (Fryer et al., 2002; Driever et al., 2009). The middle 2 cm from second leaves were soaked in 5 mM DAB solution (pH 5.0) with 0.01% Tween-20. After shaking in a dark incubator at 28°C for 2 hours, leaves were briefly dried with tissue paper, placed in a Hansatech LD2/ 3 electrode leaf chamber and subjected to 750 μmol m^-2^ s^-1^. Sampling was undertaken at 0, 5, 10, 15, 20, 30, 40, 50, 60 mins. Prior to imaging, chlorophyll was removed from the leaves by soaking in ethanol:acetic acid:glycerol (3:1:1) in a 70°C water bath for 1 hour, and then immersing in 70% (v/v) ethanol for 24 hours.

H_2_O_2_ production was also detected with 2’,7’-dichlorodihydrofluorescein diacetate (H_2_DCFDA, Sigma-Aldrich). H_2_DCFDA was dissolved in dimethyl sulfite (DMSO) to 20 mM, and then diluted with H_2_O to a final concentration of 10 μM for infiltration. Infiltration was performed as previously described (Fryer et al., 2002; Driever et al., 2009). In short, H_2_DCFDA was fed through the transpiration stream under low light (20 μmol m^-2^ s^-1^) for up to 12 hours to ensure absorption throughout the leaf. After infiltration, leaves were then clamped in the Hansatech LD2/3 electrode chamber and exposed to PPFD of 750 μmol m^-2^ s^-1^ for 0, 30, 60 minutes. Prior to confocal laser scanning microscopy, the middle vein of the leaf was imaged after generating a paradermal section by hand. To confirm whether H_2_DCFDA penetrated into the whole leaf, 30% (w/w) H_2_O_2_ solution was diluted to 100 mM with 10 μM H_2_DCFDA and used to infiltrate leaves under the same conditions as above. Leaves infiltrated with ddH_2_O at the same condition were used as controls. After infiltration, green fluorescence was assessed by confocal laser scanning microscopy.

The NADPH oxidase inhibitors diphenyliodonium (DPI) and imidazole (Sigma) were added to the DAB solution and infiltrated into leaves for 2 hours before high light treatment. DPI chloride was dissolved in DMSO to 100 mM and then diluted to 100 μM with a 5 mM DAB solution (pH 5.0, with 0.01% tween20). Imidazole was dissolved in DMSO to 1 mM and then diluted with 5 mM DAB solution (pH 5.0, with 0.01% tween20) to 20 μM. To test the effects of different CO_2_ and O_2_ contents on H_2_O_2_ products, DAB-infiltrated leaves were subjected to the high light treatment as described above, using air mixtures with different concentrations of CO_2_ controlled by an open gas exchange analyzer (LI-6400, LI-COR Biosciences, Lincoln, NE, USA) connected in line to the Hansatech LD2/ 3 electrode leaf chamber. For measurements at 2% O_2_ concentration, the LI-6400 air inlet was connected to a gas cylinder with pre-mixed 2% O_2_ in N_2_.

### Paradermal sectioning of paraffin-embedded tissue

Leaves used for paradermal sectioning had been treated with high light and chlorophyll removed. Samples were cut into small pieces of ∼ 5 mM and dehydrated in an ethanol series consisting of 70% (v/v) for 30 minutes; 85% (v/v) for 30 minutes; 95% (v/v) for 30 minutes; 100% (v/v) ethanol twice for 30 minutes, then twice in 100% (v/v) Histo-Clear II for 60 minutes each, followed by two times in paraffin wax for 60 minutes at 60°C. Melted paraffin wax was poured into 9-cm petri dishes and cut into small blocks once the wax was cool. For paradermal sectioning, blocks were trimmed so that the surface of the leaf was parallel to the surface of the block. 10 µm sections were obtained using a rotary microtome, floated in a 60°C water bath and then mounted onto clean glass slides to dry overnight in an incubator set at 42°C. Dehydration of the sections was performed as follows: 100% (v/v) Histo-Clear II twice for 10 minutes each; 100% ethanol twice for 5 minutes each; 95% ethanol (v/v); 70% (v/v) ethanol; 50% (v/v) ethanol; 30% (v/v) ethanol for 2 minutes each. Slides were then rinsed with deionized H_2_O; drained and then 30% (v/v) glycerol added prior to a coverslip.

### Laser capture microdissection

For laser capture microdissection, leaf samples were cut into ∼ 5 mm pieces and fixed in 100% (v/v) ice-cold acetone at 4°C for overnight. The next day, samples were dehydrated and embedded with Steedman’s polyester wax (Hua and Hibberd, 2019). Blocks were sectioned using a rotary microtome to 8 μm thickness and then floated on Arcturus PEN membrane slides (Fisher Scientific) with DEPC-treated water. The water was dried using tissue paper and slides were stored at −20°C and used within 12 hours after sectioning. Prior to LCM, slides were washed in 100% (v/v) ethanol for 5 mins and then air-dried for 5 mins. LCM was performed using the ArcturusXT(tm) Laser Capture Microdissection System (ThermoFisher) according to the manufacturer’s instructions. Mesophyll cells and bundle sheath stands were collected on the CapSure® Macro LCM Caps (ThermoFisher) and immediately treated with extraction buffer (from Arcturus Picopure RNA extraction kit (Thermo Fisher Scientific)) at 42°C for 30 minutes, then stored at −80°C.

### RNA extraction and quantitative PCR (qRT-PCR)

Total RNA from LCM harvested mesophyll and bundle sheath cells was extracted using Arcturus Picopure RNA extraction kit (ThermoFisher) with on-column DNaseI treatment according to the manufacturer’s protocol. cDNA synthesis for LCM samples was performed using TruSeq RNA Library Preparation Kit v2 (RS-122-2001; Illumina). Total RNA from whole leaves was extracted from 100 to 200 mg of fully expanded leaves using Triazol reagent (Sigma-Aldrich) according to the manufacturer’s instructions. Superscript II reverse transcriptase (18064-022; ThermoFisher Scientific) was used for cDNA synthesis of whole leaves. qRT–PCR was performed with the Bio-Rad CFX384 Real-Time PCR system using the SYBR Green Jumpstart Taq ReadyMix (S4438-100RXN; Sigma-Aldrich) with the following PCR conditions: 94°C for 5 mins and 40 cycles of 94°C for 10 s, 60°C for 1 min. Relative gene expression levels were calculated using the 2^-ΔΔ*CT*^ method (Kubista et al., 2006) and normalized with *OsUBQ5*. Gene-specific primer sequences are listed in Supplemental Table 2. For each gene, three biological and three technical replicates were performed.

### RNA-sequencing library preparation and data processing

RNA integrity and quality of LCM samples were assessed using 2100-Bioanalyzer (Agilent Technologies, USA) with an Agilent Bioanalyser RNA 6000 Pico assay and QuBit (Thermo Fisher Scientific), respectively. Only samples RNA Integrity Number (RIN) ≥4.4 were selected for the final sample cohort. 50∼150 ng starting total RNA from 12∼15 paradermal sections of each replicate were used for RNA-seq library construction using the QuantSeq 3’ mRNA-Seq Library Prep Kit (Lexogen) according to the manufacturer’s recommendations. cDNA libraries were assessed using 2100-Bioanalyzer (Agilent Technologies, USA) before 100 bp single-end sequencing using NextSeq500 (Illumina) system at the Department of Biochemistry Sequencing Services at the University of Cambridge based on standard protocols. Three biological replicates were conducted for each cell type. Data processing was performed using custom scripts. Briefly, raw reads were processed using Trimmomatic, mapped to the reference rice transcriptome genome (MSU7.0, http://rice.plantbiology.msu.edu/index.shtml) and performed quantified using Salmon (Patro et al., 2017). Differential expression analysis was performed using DESeq2. Stringent criteria with log_2_ fold change (log_2_FC) >0.5 and adjusted p value (padj) < 0.05 were used to screen the differentially expressed genes (DEGs) between bundle sheath strand and mesophyll cells. Plots were generated with custom scripts in RStudio using the package ggplot2. Three biological replicates for each cell type were performed.

### Generation of *OsRBOHB*-overexpression (*OsRBOHB*-OE) lines

To generate the *OsRBOHB*-OE plants, the full-length coding region of *OsRBOHB* was was amplified from the first-strand cDNA of Nipponbare and insert into vector pCAMBIA1301 under the control of ubiquitin promoter. The resulted construct was transformed into Nipponbare by Agrobacterium-mediated transformation. The primers used for the construct are listed in Supplemental Table 2.

### Imaging

Intact leaves were imaged with Leica m165 FC microscopy. The paradermal sections were imaged with Olympus BX41 microscopy. DAB staining intensity was quantified using ImageJ (FIJI build, version 1.52q, NIH, USA). Confocal micrographs for detecting the DCF fluorescence were taken using a Leica SP8 confocal microscopy (excitation 488 nm, barrier 515–555 nm).

### Statistical analysis

All statistical analyses were conducted in R (v.3.6.3). Two-way analysis of variance (ANOVA) was used to assess statistical differences in DAB staining between BSS and M cells after a time course of high light treatment, and the effects of different concentrations of CO_2_ and O_2_ concentrations on H_2_O_2_ production under high light treatment. Three-way ANOVA was used for statistical analysis on the effects of different concentrations of CO_2_ and O_2_ concentrations on H_2_O_2_ production under high light treatment and also used for assess the data in experiments with the inhibitors DPI, imidazole, and also used to compare the H_2_O_2_ production of *OsRBOHA*-OE and its wildtype under a time course of high light treatment. Tukey Honest Significant Differences (TukeyHSD) test was performed for multiple pairwise-comparison between the means of groups. Levene’s test was used to check the homogeneity of variances and Shapiro-Wilk test was used to check the normality assumption. Unpaired T-tests were performed to compare differences in transcript abundance between BSS and M cells and also used to compare the differences in DAB staining between wild type and *OsRBOHB*-overexpression lines and mutants under dark/high light conditions.

## Acknowledgements

We thank Ivan Reyna-Llorens (University of Cambridge) for making the phylogenetic tree. The work was funded by China International Postdoctoral Exchange Fellowship to HX, ERC grant RG80867 Revolution to JMH and a C_4_ Rice project grant from The Bill and Melinda Gates Foundation to the University of Oxford (2015–2019).

---

**Figure S1.**
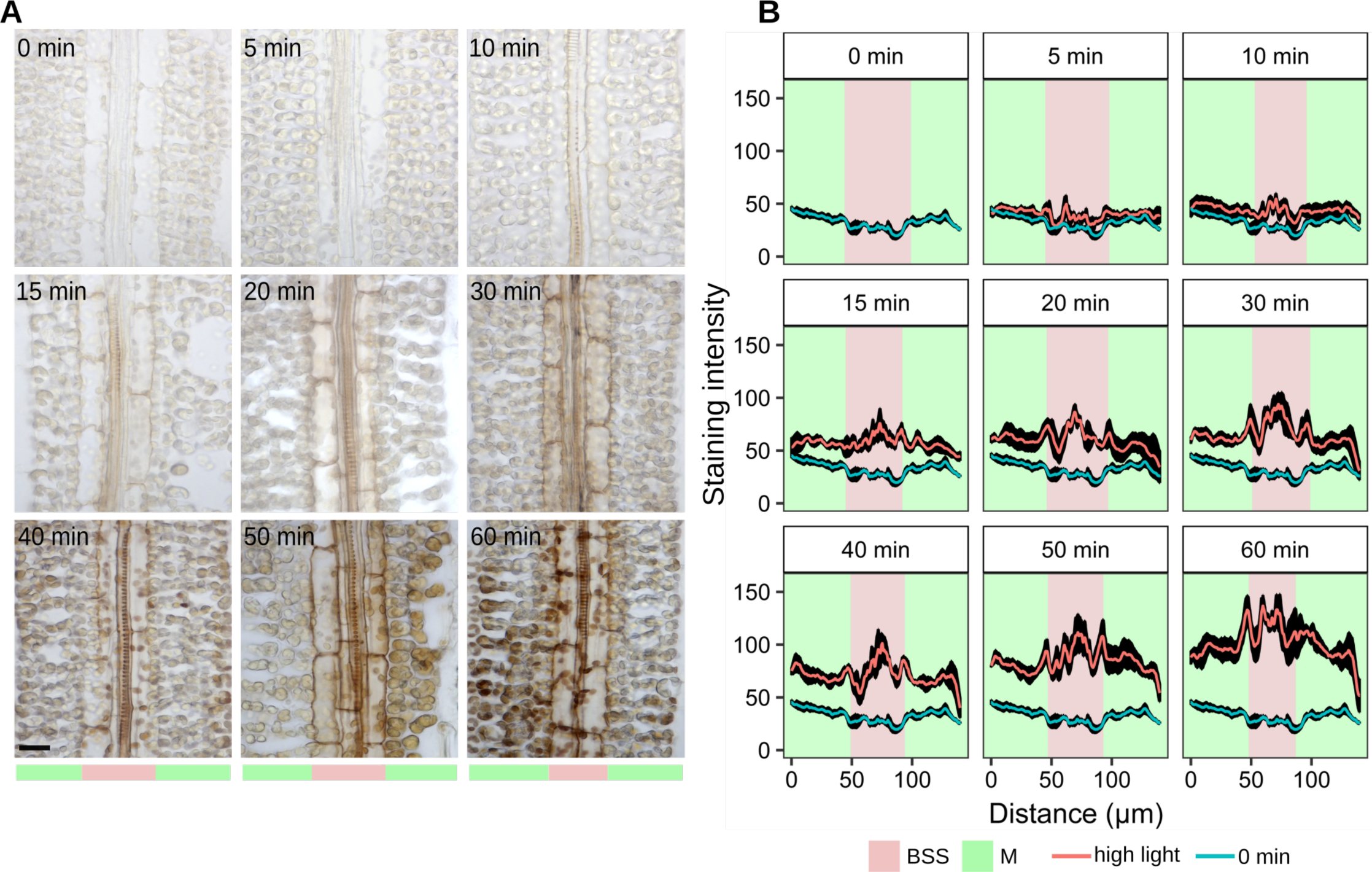
DAB staining of paradermal sections from rice leaves exposed to high light. (A) Representative images of leaves exposed to 750 μmol m^-2^ s^-1^ photon flux density for 0, 5, 10, 15, 20, 30, 40, 50 or 60 minutes. Brown stain indicates the formation of the polymerization product formed in the presence of reactive oxygen species. Scale bar denotes 10 μm. (B) Quantitation (arbitrary units) of DAB staining from paradermal sections illustrating the extent to which reactive oxygen species are first detected in vein and bundle sheath strands versus mesophyll cells The x-axis represents a scan from left to right across the paradermal sections. The y-axis depicts gray values extracted from paradermal sections. Data are presented as mean (red or blue line) and one standard error from the mean, n = 4). The response of bundle sheath strands to high light was greater than that of mesophyll cells (Two-way ANOVA p < 0.001).

**Figure S2.**
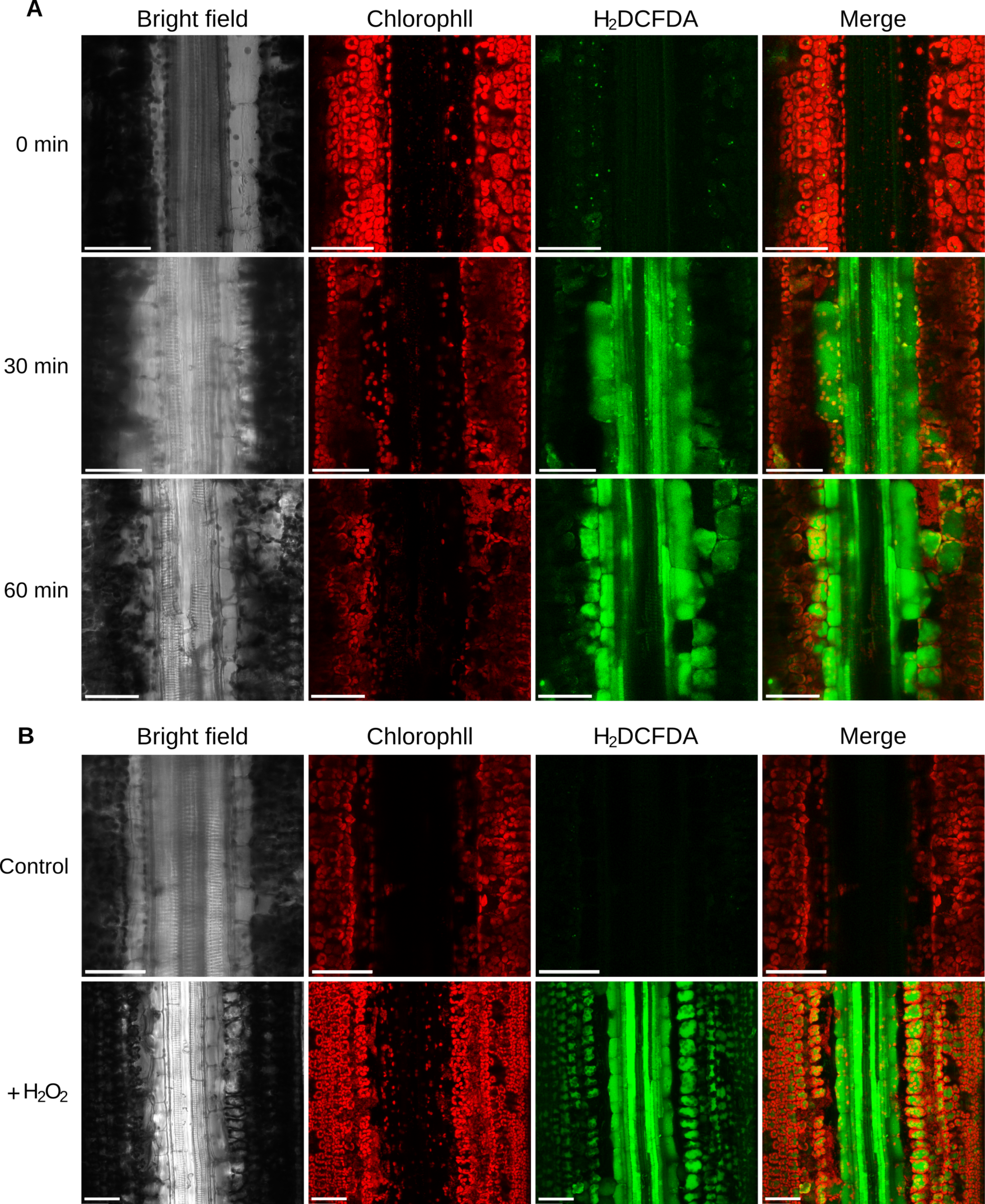
Confocal scanning microscopy indicates bundle sheath preferential fluorescence from H_2_DCFDA in rice leaves exposed to high light. (A) Representative images of leaves were fed with 10 μM H_2_DCFDA for 12 h prior to being exposed to high light (750 μmol m^-2^ s^-1^ photon flux density) for the times indicated. (B) Representative images from control leaves suppled with water as well as those suppled with 10 μM H_2_DCFDA and exogenous 100 mM H_2_O_2_. Fluorescence from H_2_DCFDA was not detected in control leaves, but in those to which H_2_O_2_ had been added it was detected in both mesophyll and bundle sheath strands. Scale bars denote 50 μm.

**Figure S3.**
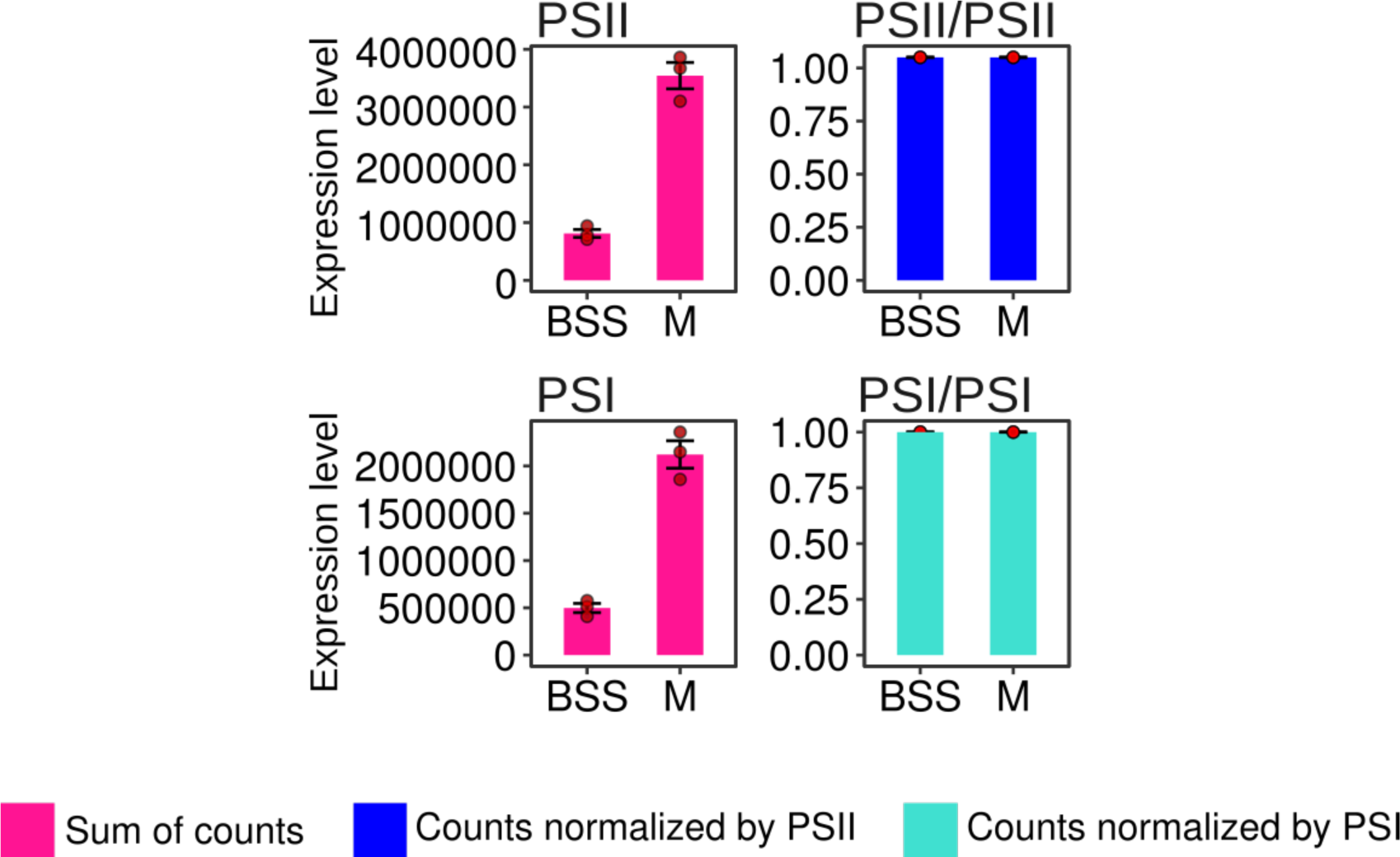
Normalisation of PSII and PSI against themselves indicates the expected one-to-one relationship.

**Figure S4.**
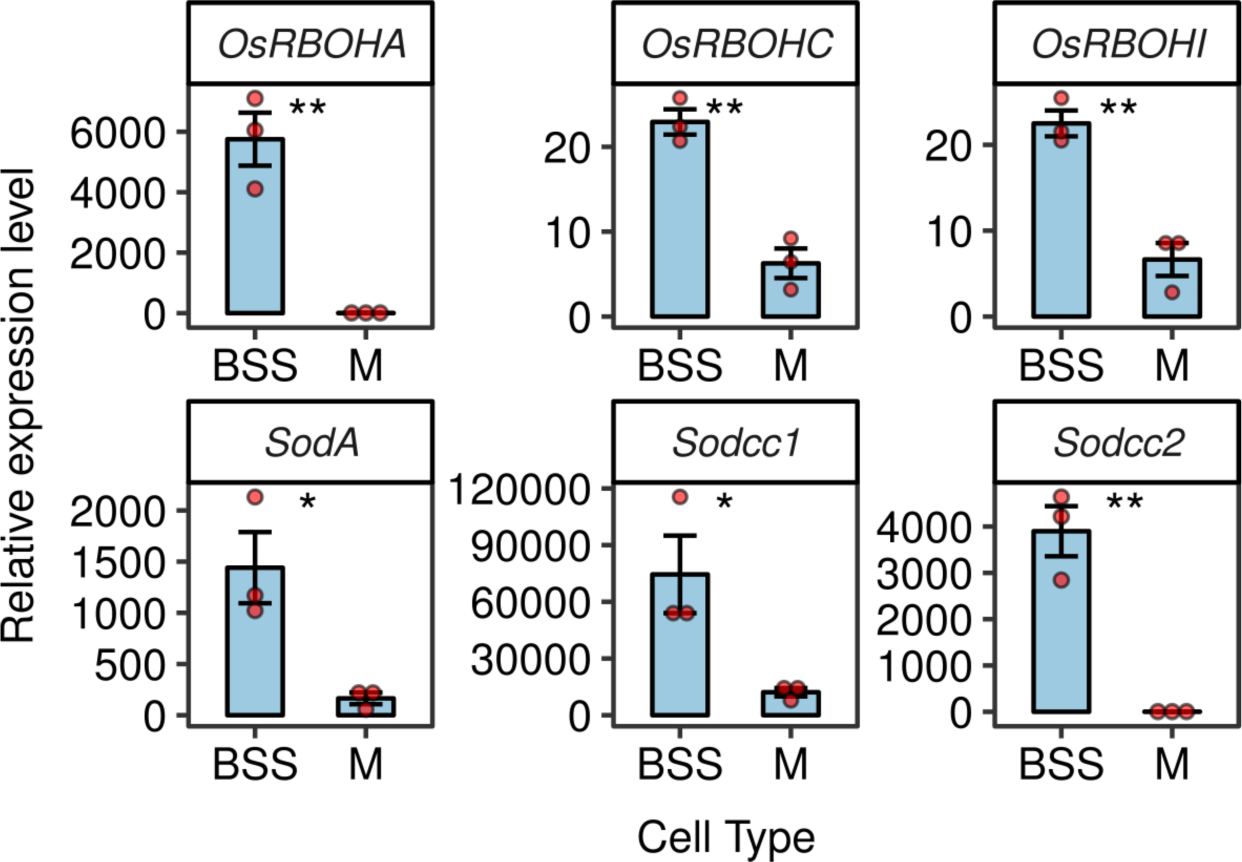
qPCR confirms analysis from deep sequencing and shows that transcripts of *RBOHA, RBOHC & RBOHI* as well as *SODA, SODCC1 & SODCC2* are more abundant in bundle sheath stands (BSS) than mesophyll (M) cells. RNA extracted from green leaves. Data are normalized to *OsUBQ5* as a control and presented as mean ± one standard error, n = 3. T-tests indicate statistically significant differences (** p< 0.01, * p< 0.05).

**Figure S5.**
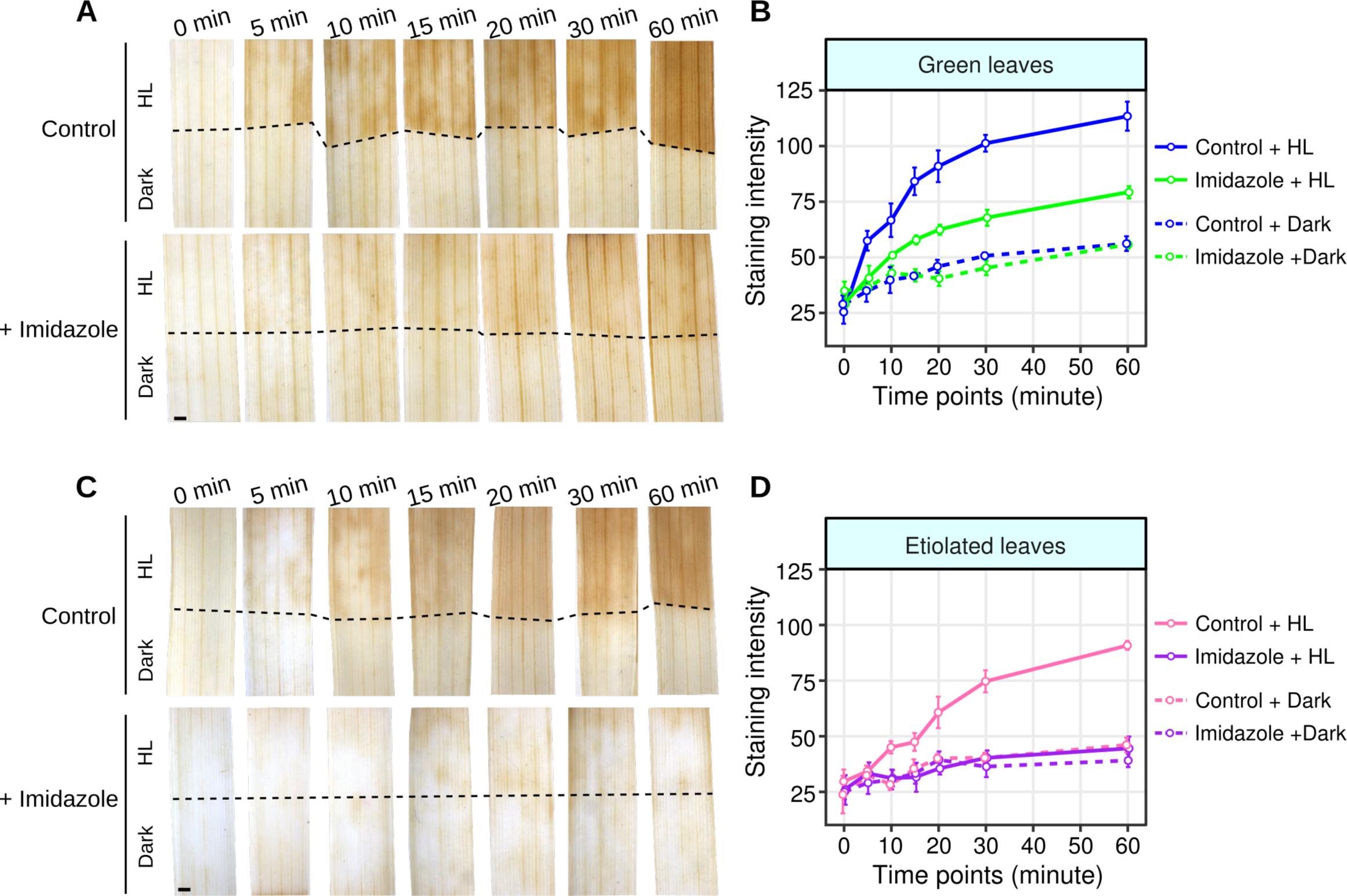
Inhibiting NAD(P)H oxidase activity with imidazole phenocopies the effect from diphenyleneiodonium in green and etiolated leaves. (A&B) Imidazole partially inhibits the accumulation of DAB in green leaves. (C&D) Imidazole completely inhibits DAB accumulation in etiolated leaves. ANOVA showed DAB staining was reduced after Imidazole treatment in green leaves (p<0.001), etiolated leaves (p<0.001).

**Figure S6.**
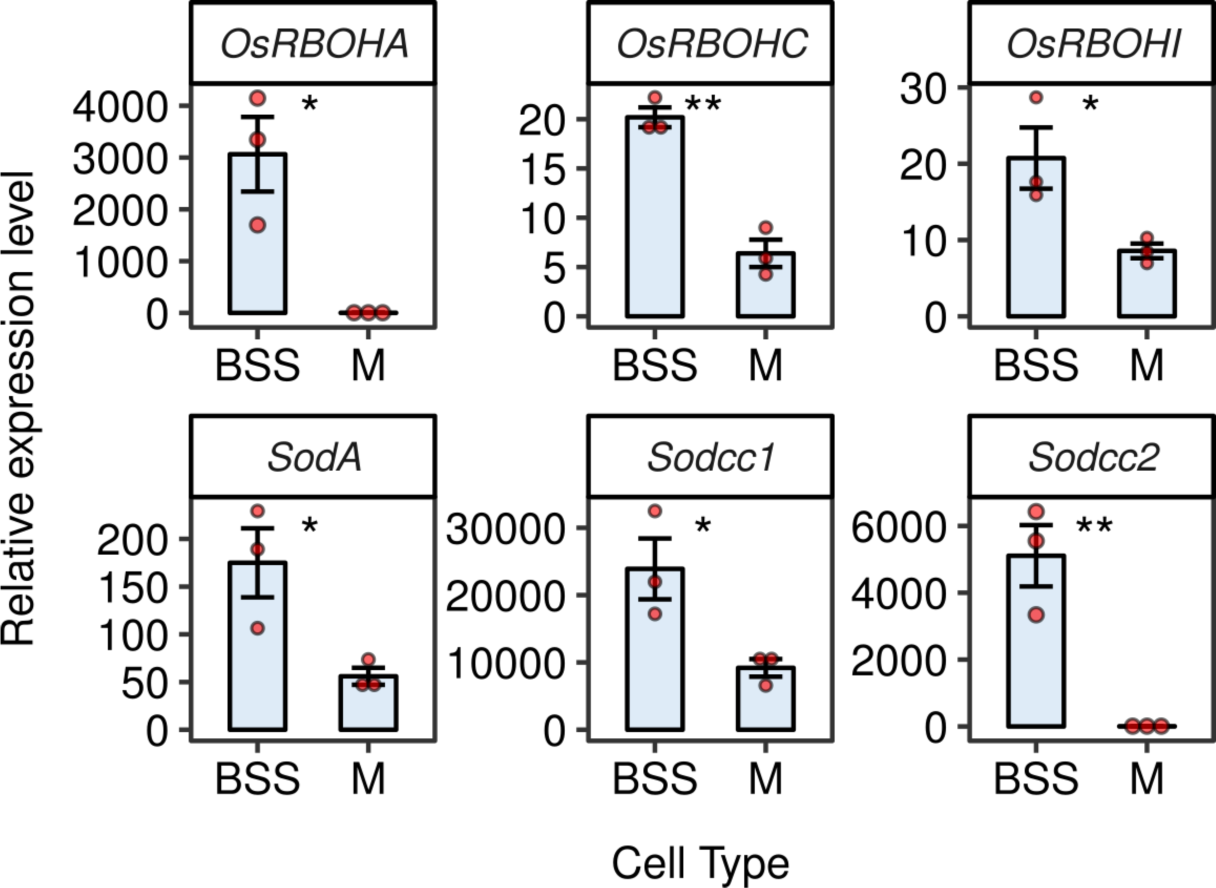
Q-PCR shows that transcripts encoding *OsRBOHA, RBOHC & RBPOHI* as well as *SODA, SODCC1 &SODCC2* are more abundant in bundle sheath strands than mesophyll ells in etiolated leaves of rice. *OsUBQ5* was used to normalize expression of each gene, and data are presented as the mean ± one standard error, n = 3. T-tests showed statistically significant difference between bundle sheath strands (BSS) and mesophyll (M) cells with ** = p< 0.01, * = p < 0.05.

**Figure S7.**
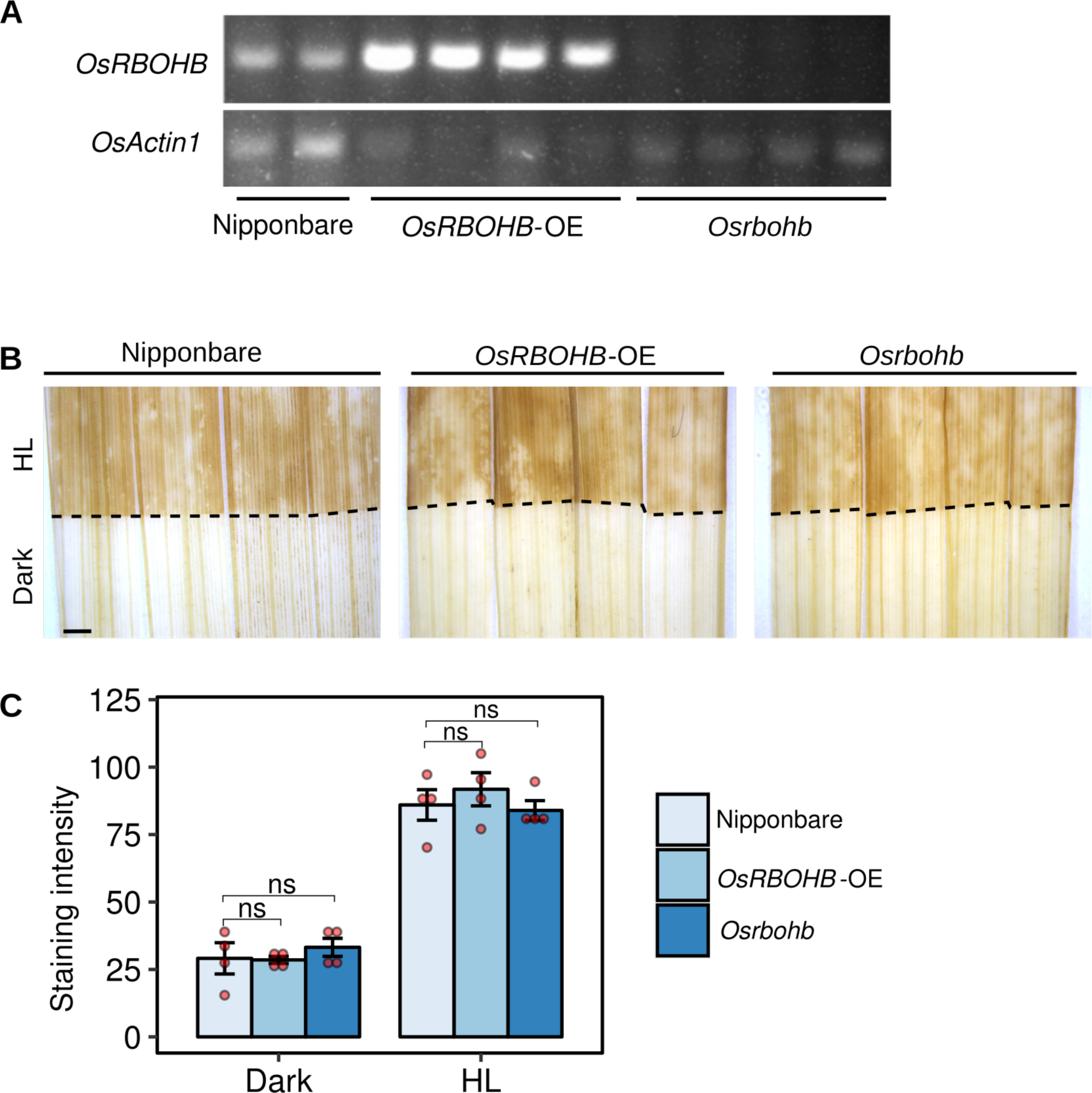
The *RBOHB* gene appears not to be involved in the high light response of the rice bundle sheath. (A) Semi-quantitative RT-PCR indicating that compared with the Nipponbare control, an over-expression line (*O*s*RBOHB-OE*) and a mutant allele (*Osrbohb*) contain increased and reduced transcripts from *RBOHB* respectively. *ACTIN1* is shown below as a control. (B) Representative images showing that the over-expression line and mutant allele of *RBOHB* show no difference in DAB staining compared with the Nipponbare control. (C) Quantification of DAB staining in (B). Scale bar represents 2 mm. T-tests showed that there was no significant (ns) statistical differences associated with over-expression of *OsRBOHB*.

**Figure S8.**
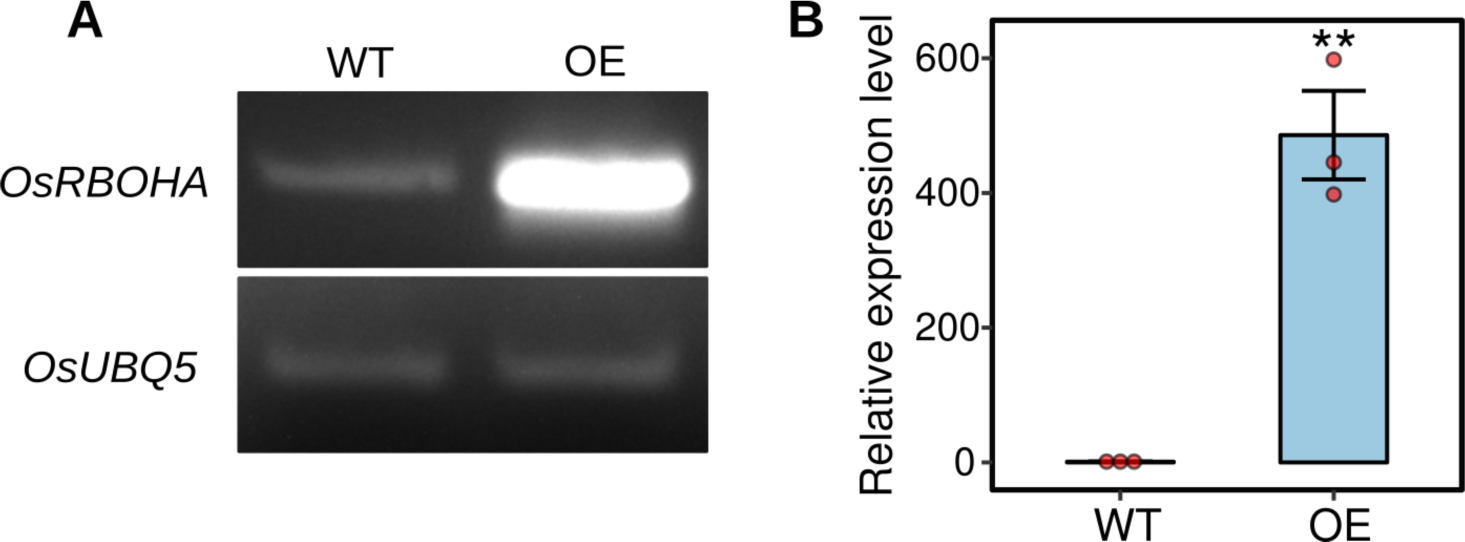
RT-PCR (A) and Q-PCR (B) were used to confirm over-expression of *OsRBOHA* compared with wild type (WT). *OsUBQ5* was used as a control. T-test showed statistically significant difference in the over-expressor (OE) compared with wild type (WT) (p<0.01).

**Figure S9.**
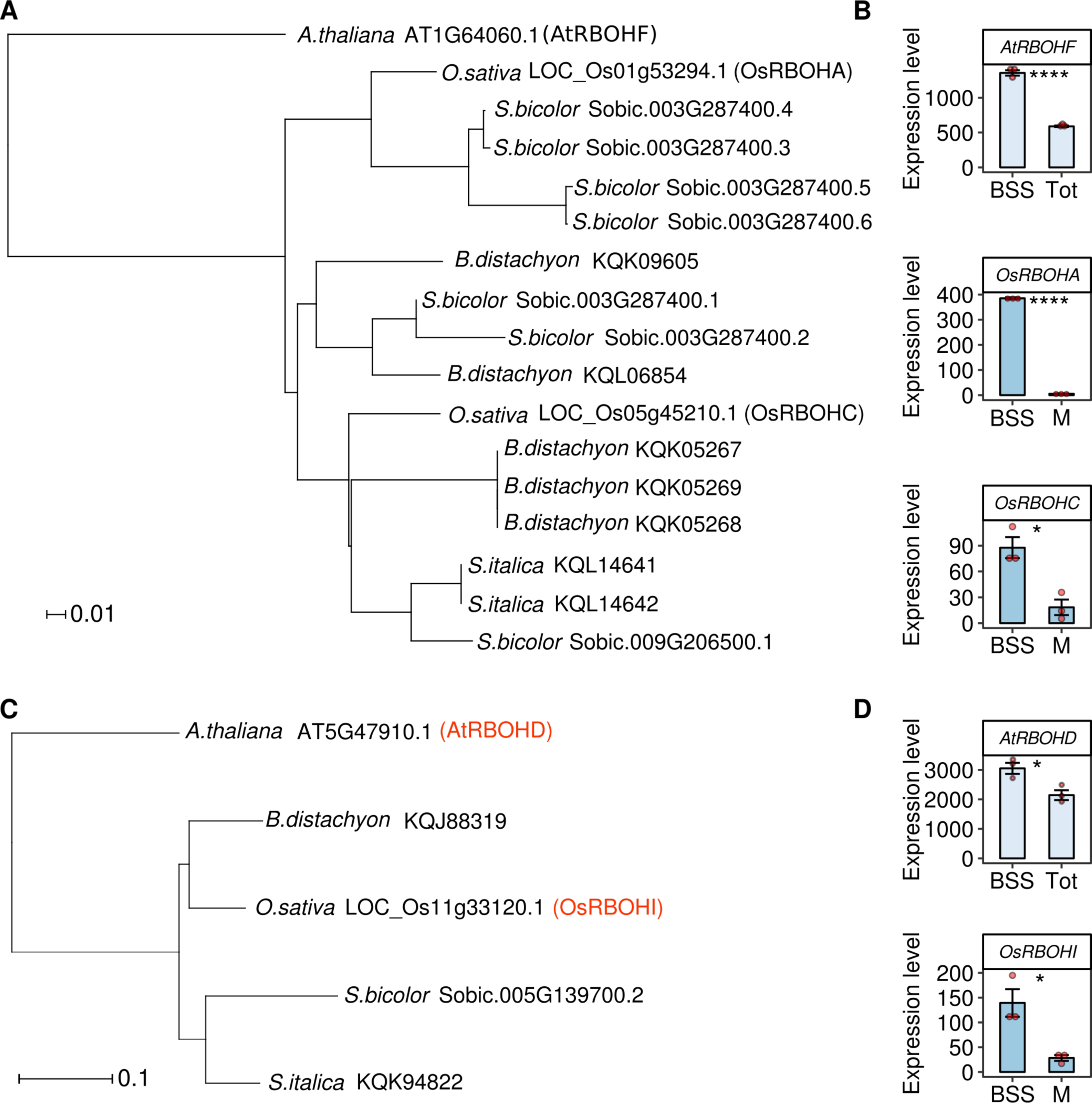
Phylogenetic reconstructions of *RBOH* genes depicting orthology relationships between those from rice, *Arabidopsis thaliana, Sorghum bicolor*, and *Brachypodium distachyon*. (A) Phylogenetic reconstruction indicating likely relationship between *AtRBOHF* and related genes in the grasses. (B) Transcript abundance of *AtRBOHF* in bundle sheath strands and total leaf material of *A. thaliana*, and of *OsRBOHA* and *OsRBOHC* in bundle sheath strands and mesophyll cells of rice. (C) Phylogenetic reconstruction indicating likely relationship between *AtRBOHD* and related genes in the grasses. *AtRBOHD* and *OsRBOHI* are highlighted in red. (D) Transcript abundance of *AtRBOHD* in bundle sheath strands and total leaf material or *A. thaliana* and of *OsRBOHI* in bundle sheath strands and mesophyll cells of rice. T-tests show statistically significant differences between cell type **** = p < 0.0001, * = p < 0.05.

